# Towards a broad-spectrum antiviral, the myristoyltransferase inhibitor IMP-1088 suppresses viral replication – the Yellow fever NS5 is myristoylated

**DOI:** 10.1101/2021.03.09.434547

**Authors:** Melissa Immerheiser, Melissa Zimniak, Helen Hilpert, Nina Geiger, Eva-Maria König, Jochen Bodem

**Affiliations:** Institut für Virologie und Immunbiologie, Universität Würzburg, Versbacher Str. 7, 97078 Würzburg, Germany

**Keywords:** Dengue virus, Yellow fever virus, HIV-1, IMP-1088, antiviral, NS5

## Abstract

Although a potent Yellow fever vaccine is available since 1937, up to 200.000 severe cases are reported per year, which indicates that virus vaccines require additional support by antiviral therapies. Direct-acting antiviral drugs against severe and widespread diseases, such as DENV and Yellow fever infections with more than millions of diagnosed diseases per year, are still not available. Since antivirals’ development against neglected diseases is uneconomical, a broadspectrum antiviral compound would be of public benefit. Here, we show that IMP-1088, a recently published myristoyltransferase-1/2 inhibitor suppressing Rhino- and Polioviruses, inhibits replication of HIV-1, Yellow fever virus, Dengue virus, Vaccinia virus, CMV, and human Herpesvirus 8 in the low nanomolar range, indicating that IMP-1088 has broad-range activity against different pathogenic virus families. The inhibition relies on virally encoded myristoylation signals since Zika, Chikungunya, and Enterovirus 71 are not affected by IMP-1088. Furthermore, we show that the Yellow fever NS5 protein is myristoylated and IMP-1088 treatment of Dengue and Yellow fever infected cells leads to a re-localisation of the viral NS5 proteins.

**Author Summary:** Treatment of viral diseases requires the development of tailored drugs specific to inhibit certain virus families. This specificity results in missing treatment options for important human pathogens such as Yellow fever and Dengue virus infection since the development is laborious and costly. Substances acting on various virus families could solve this problem. Here, we describe that IMP-1088, an inhibitor of the cellular myristoyltransferase, inhibits HIV-1, Dengue virus, Yellow fever viruses, Vaccinia virus, and Herpesviruses at low concentrations, which do not affect cell proliferation. Viruses without predicated myristoylation sites, such as Zika viruses, were not inhibited by IMP-1088. Since no experimental evidence was provided that Yellow fever virus proteins are myristoylated, we analysed the post-translational modification of Yellow fever NS5 protein. We determined the subcellular localisation to understand the mechanism of the IMP-1088 mediated suppression and could show that both the Dengue and the Yellow fever NS5 proteins are re-localised by IMP-1088 treatment.

## Introduction

During the last decades, fundamental progress in treating acute and chronic viral infections has been made. Antivirals for the treatment of Hepatitis C (1), B (2) and D (3) viruses, the human immunodeficiency virus (HIV-1) (4), Herpesviruses, and Cytomegalovirus (CMV) have been developed and improved. Despite this success, neglected tropical viral diseases such as Dengue virus (DENV) infections resulted in significant outbreaks causing millions of manifest clinical disorders in the endemic regions without the possibility to treat infected patients. The number of DENV infections increased by more than 150%, from 2.2 million cases in 2010 to 3.34 million in 2016. Even in Germany, where the virus is not endemic, imported DENV cases in 2017 and 2018 surpassed the endemic tick-borne encephalitis cases. In 2015 a DENV vaccine was approved, but it is only recommended to seropositive individuals since it may increase the risk for severe DENV infections in seronegative persons.

However, the development of effective vaccines against Chikungunya and other mosquito-borne diseases might improve the situation. Generally, it is believed that vaccines might finally eliminate DENV and other tropical infections. Still, the global status of Yellow fever virus (YF) infections, where a live-attenuated vaccine is available at low costs since 1937, might shed another light on this assumption. In the year 2018, in a recent YF virus outbreak in Brazil, 237 deaths occurred, and still, 200.000 humans are infected, with 30.000 deaths each year (5, 6). Thus, effective antiviral therapy is still urgently needed.

Most antiviral drugs used to treat viral infections target viral proteins by inhibiting either virus entry or viral enzymes, such as polymerases, integrase or protease. These treatment strategies can result in rapid viral adaptation leading to the development of viral resistance. It was recently published that IMP-1088, an inhibitor of the cellular myristoyltransferase, will suppress Polio-and Rhinoviruses’ replication (7). Interestingly, myristoylated viral proteins are involved in the replication of several human pathogenic viruses at different stages of the viral life-cycle. While Vaccina L1R and A16L proteins of the fusion complex were shown to be myristoylated, myristoylation of the HIV-1 Gag protein is essential for assembly. The requirement of post-transcriptional myristoylation for the replication of RNA and DNA viruses offers the rare possibility to use a single drug active against different and unrelated viruses. Furthermore, since this inhibitor acts on a cellular enzyme, viral adaption seems to be more unlikely.

This report presents evidence that IMP-1088 inhibits potently, specifically, and selectively replication of HIV-1, Vaccinia virus, human Herpesvirus 8 (HHV-8) and mouse CMV. Furthermore, we show that the YF NS5 protein is myristoylated and that the recently described substance suppresses the YF and DENV. We provide evidence that this effect is virus-specific since the replication of other viruses, such as Zika, Chikungunya or Rubella viruses, are not affected by IMP-1088.

## Results and Discussion

### IMP-1088 inhibits HIV-1 replication

Myristoylation of the HIV-1 Gag and Nef proteins is essential for virus assembly at the plasma membrane and the function of Nef. Thus, HIV-1 can serve as proof of principles, whether the inhibition by myristoyltransferase influences viral replication. We isolated PBMCs and treated the cells with IMP-1088 at concentrations of 10nM and 100nM. These concentration were non-toxic in MTT assays. The cells were subsequently infected with NL4-3 derived virus in triplicates to analyse whether IMP-1088 inhibits HIV-1 replication. Cellular supernatants were collected three days after infection, and viral infectivity was determined using TZM-bl indicator cells (Figure 1). IMP-1088 inhibited viral infectivity by two orders of magnitude at concentrations of 100nM, indicating that suppressing the myristoyltransferase activity decreased viral infectivity. Similar results were obtained with MT4 cells. These results open the possibility for new treatment options for patients with highly resistant HIV-1 viruses. IMP-1088 did not show any toxic effect on PBMCs at the concentrations used in these experiments.

**Figure 1.**
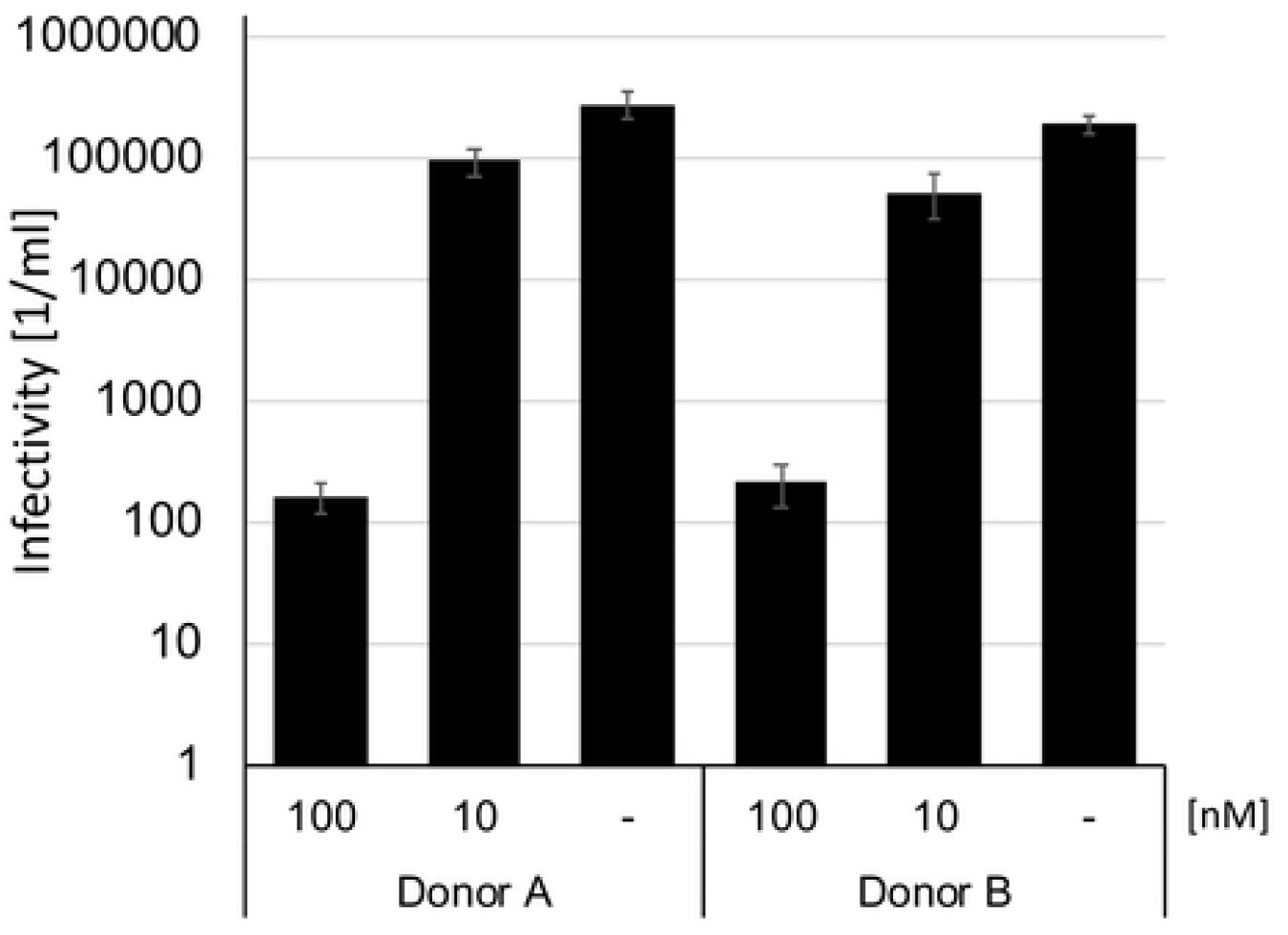
IMP-1088 suppresses HIV-1 replication. PBMCs from two different donors were treated with IMP-1088 and subsequently infected with HIV-1 in triplicate assays. Viral supernatants were collected three days after infection, centrifuged to remove detached cells and titrated on TZM-bl indicator cells. Infected cells were visualised by beta-galactosidase stain, and viral titers were calculated.

### Yellow fever NS5 is myristoylated

The plus-stranded RNA viruses like DENV, Poliovirus, and YF encode a single open reading frame, which is structured in the coding regions of the structural proteins and the coding regions of the non-structural (NS) proteins. In Polioviruses, the start codon used to translate this precursor protein is followed by a glycine residue, which is myristoylated and finally leads to a modified VP4 protein. The DENV and YF precursor proteins lack this myristoylation site. Cellular proteases and the viral protease are responsible for cleaving the viral precursor polyprotein into functional proteins. The viral precursor protein processing is essential for viral infectivity and results in functional subunits with predicted myristoylation sites in the NS5 protein (8). Thus, IMP-1088 might inhibit the viral replication of other viruses too.

It has recently been suggested that the NS5 proteins of DENV and YF could be myristoylated, but neither the NS5 myristoylation nor its influences have been determined so far. Thus, we analysed whether the amino terminus of the YF proteins contains a potential myristoylation site using bioinformatics (9), and the N-terminal amino acid sequence of NS5 was predicted representing myristoylated signals with an ExPASy score of 0.935. This led to the hypothesis that the Yellow fever precursor protein could be cleaved by viral and cellular proteases, and subsequently, the NS5 protein could be myristoylated. HEK293T cells were transfected with a codon-optimised NS5-expression plasmid (pCDNA-3.4-YFsynNS5-FLAG), encoding a FLAG-epitope at the carboxyl terminus to demonstrate that NS5 is myristoylated. First, the NS5 expression was visualised with an anti-FLAG-antiserum by Western blotting. Next, the transfected cells were lysed, and NS5 was affinity purified with M2-Flag-Resin. The eluted protein was visualised on the coomassie stained PAGE and excised. The protein was digested with either Elastase, Thermolysin, Trypsin or Trypsin+10%Acetonitril to obtain overlapping peptides, which were subjected to mass spectrometry. We found that the glycine residue at the NS5 cleavage site is myristoylated or acetylated (Figure 2). These results indicated that YF might be sensible to IMP-1088 as well.

**Figure 2.**
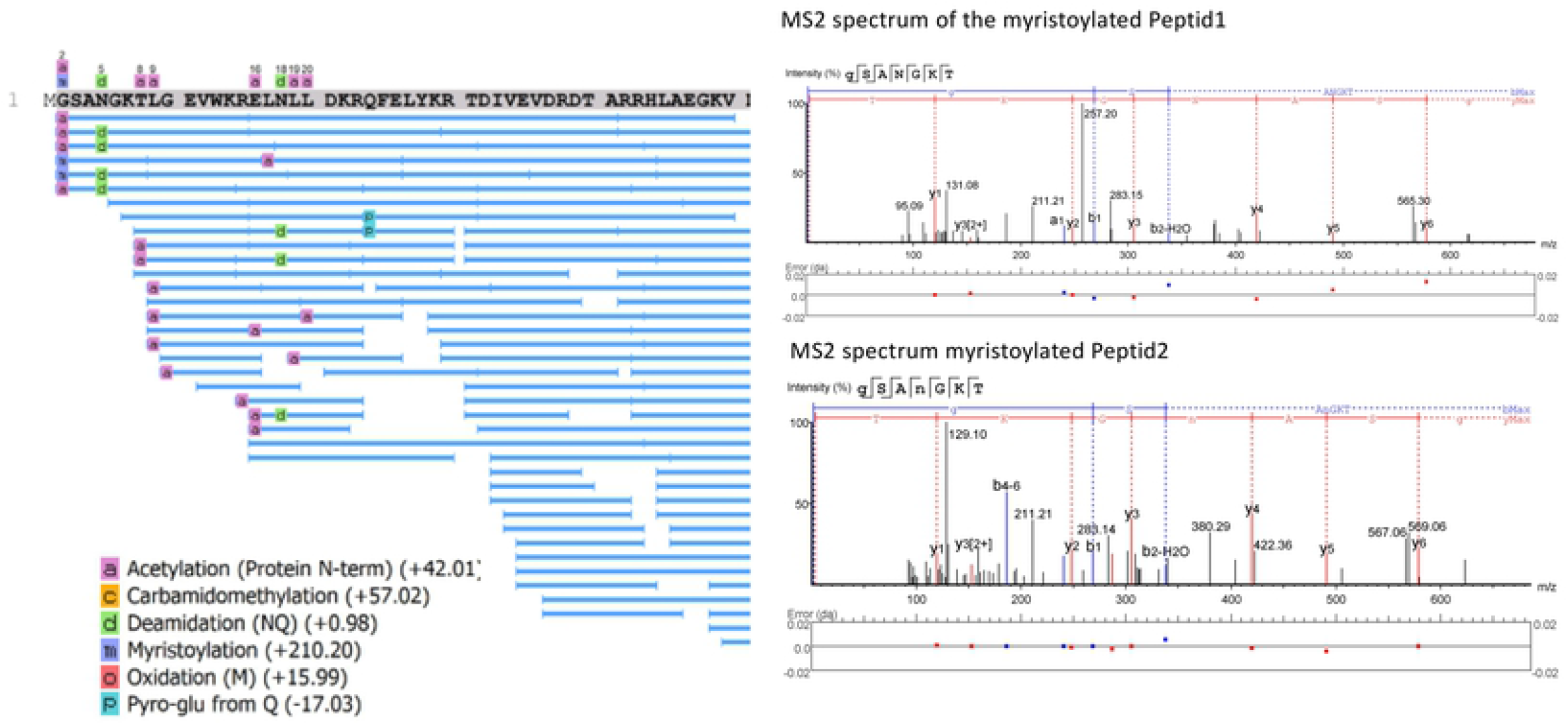
The Yellow fever virus NS5 is myristoylated. HEK293T cells were transfected with a codon-optimised Yellow fever NS5-FLAG expression plasmid. The NS5 proteins were affinity purified with anti-FLAG agarose beads and separated on SDS-PAGE. The eluted protein was digested with Elastase, Thermolysine, Trypsin or Trypsin +10%Acetonitril to generate overlapping peptides. The protein modifications were analysed by mass spectrometry. Depicted are the analysed peptides and the tandem mass spectra of the myristoylated petides.

### IMP-1088 suppresses YF replication

Before analysing the effects of IMP-1088 on the YF replication, the influence of the compound on cellular growth was determined. Vero cells were seeded in optical 96 well plates. On the next day, cell numbers per individual well were determined. Afterwards, cells were incubated with 500nM, 100nM, or 10nM of IMP-1088 for two days, and again all cells per well were counted. We did not observe any influence of IMP-1088 on cellular growth. Next, Vero cells were incubated with either IMP-1088 or DMSO as control and subsequently infected with the YF. Cellular supernatants were collected two days after infection, viral RNAs were isolated, and RTqPCR determined genome copy numbers with primers and probes described before (10). The efficiency of the RTqPCR was determined with RNA dilutions rows. Viral copy numbers were quantified with a synthetic standard of known concentration. IMP-1088 decreased the number of viral genome copies at 100nM and 10nM by two or one orders of magnitude, respectively (Figure 3), indicating that IMP-1088 is a potent inhibitor of YF replication. The EC50 was determined with 11.3 ± 1.9 nM.

**Figure 3.**
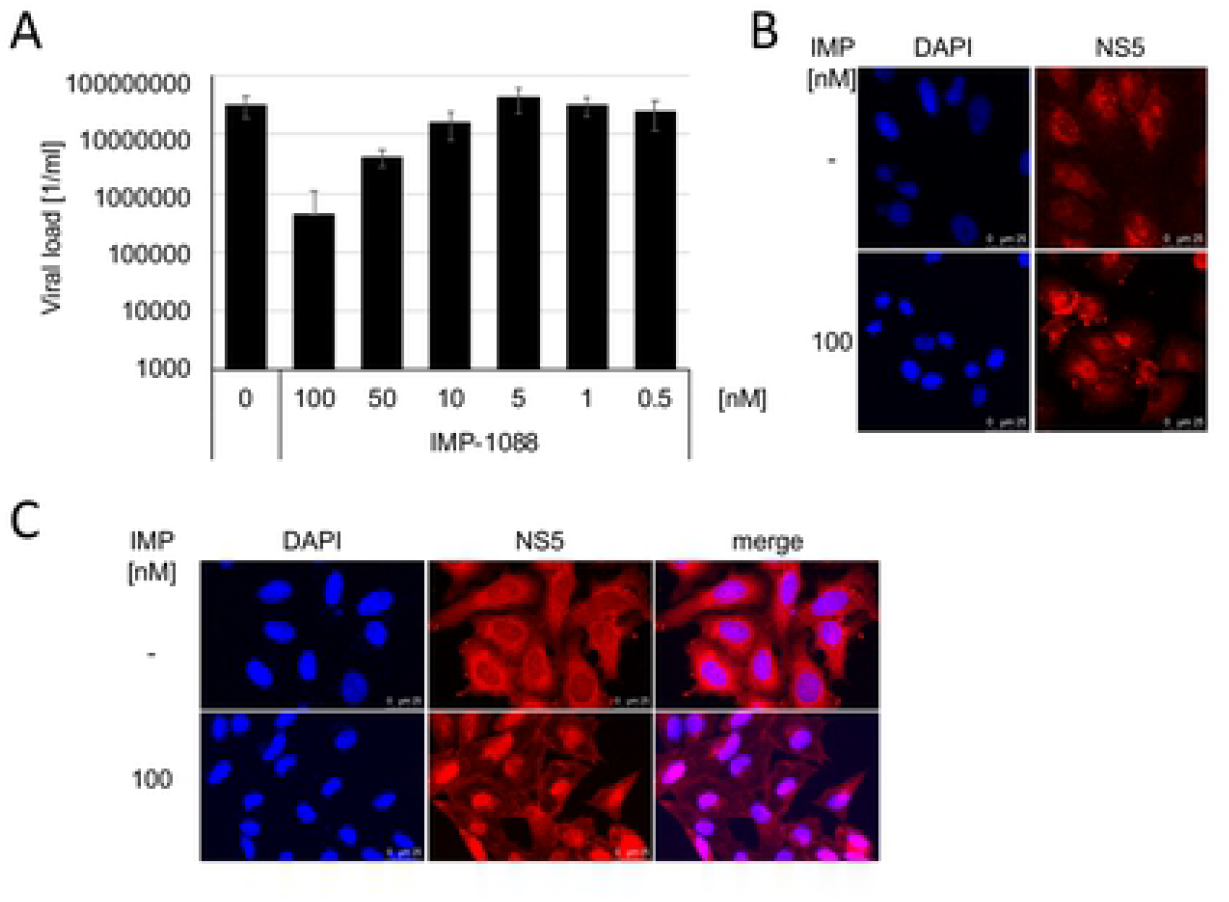
IMP-1088 inhibits Yellow fever virus replication and changes the subcellular localisation of NS5. **(A)** Vero cells were incubated with IMP-1088 and infected with the Yellow fever virus in triplicates. Released viruses were determined by RTqPCR. The EC50 was calculated. Bars indicate the mean of three independent infections. The error bars represent the standard deviation. **IMP-1088 changes the subcellular localisation of the Yellow fever NS5 protein**. HepG2 cells were infected with the Yellow fever virus **(B)** or transfected with the FLAG-tagged codon-optimised NS5 expression plasmid **(C)**. The subcellular localisation of NS5 was determined by immunofluorescence with an NS5 specific antibody (B) or the M2-FLAG antiserum (C).

### IMP-1088 changes the subcellular localisation of YF NS5

Since myristoylation of NS5 might influence membrane targeting and, thereby, the subcellular localisation of the protein, microscopic analyses with an NS5-specific antibody were performed. Vero cells were seeded on microscopic slides, infected with YF, and fixed with paraformaldehyde two days after infection. Staining with the NS5 specific antibody revealed that YF NS5 was localised in the cytoplasm and nucleus in infected Vero cells. In contrast, NS5 in YF infected cells treated with 100nM IMP-1088 was localised almost exclusively in the nucleus (Figure 3). This indicates that the cytoplasmic localisation of the YF NS5 protein is myristoylation dependent. In order to show that no other viral protein is required for the IMP-1088 dependent re-localisation, a FLAG-tagged codon-optimised NS5 was synthesised. HepG2 cells were seeded on microscopic slides and transfected with pcDNA3.4-synNS5-FLAG. Cells were fixed and permeabilised two days after transfection. The subcellular localisation was visualised with a rabbit-anti-FLAG antibody (Figure 3). Again, the NS5 protein was localised in the cytoplasm and nucleus. Treatment with IMP-1088 resulted in re-localisation of the NS5 protein to the nucleus showing that no other viral protein is required for the change in subcellular localisation. These results provide evidence that the NS5 localisation is dependent on the myristoylation. Furthermore, the non-myristoylated form of NS5 seems to be localised almost exclusively in the nucleus.

### IMP-1088 suppresses Dengue viruses specifically and leads to re-localisation of NS5

Since YF responded so well to IMP-1088, we sought to analyse other plus-stranded RNA viruses, such as Zika and Dengue. In similar experiments, Vero cells were incubated with IMP-1088 and infected with Dengue virus serotype 2 (DENV2). After 3 days, viral RNAs were isolated and quantified by RTqPCR. 100nM IMP-1088 suppressed viral loads about one order of magnitude, while 10nM of IMP-1088 reduced viral genome amounts to 45%. This indicated that DENV2 is susceptible to IMP-1088 (Figure 4). In addition, the EC50 values for DENV2 (0.15 ± 0.14 nM) were calculated.

**Figure 4.**
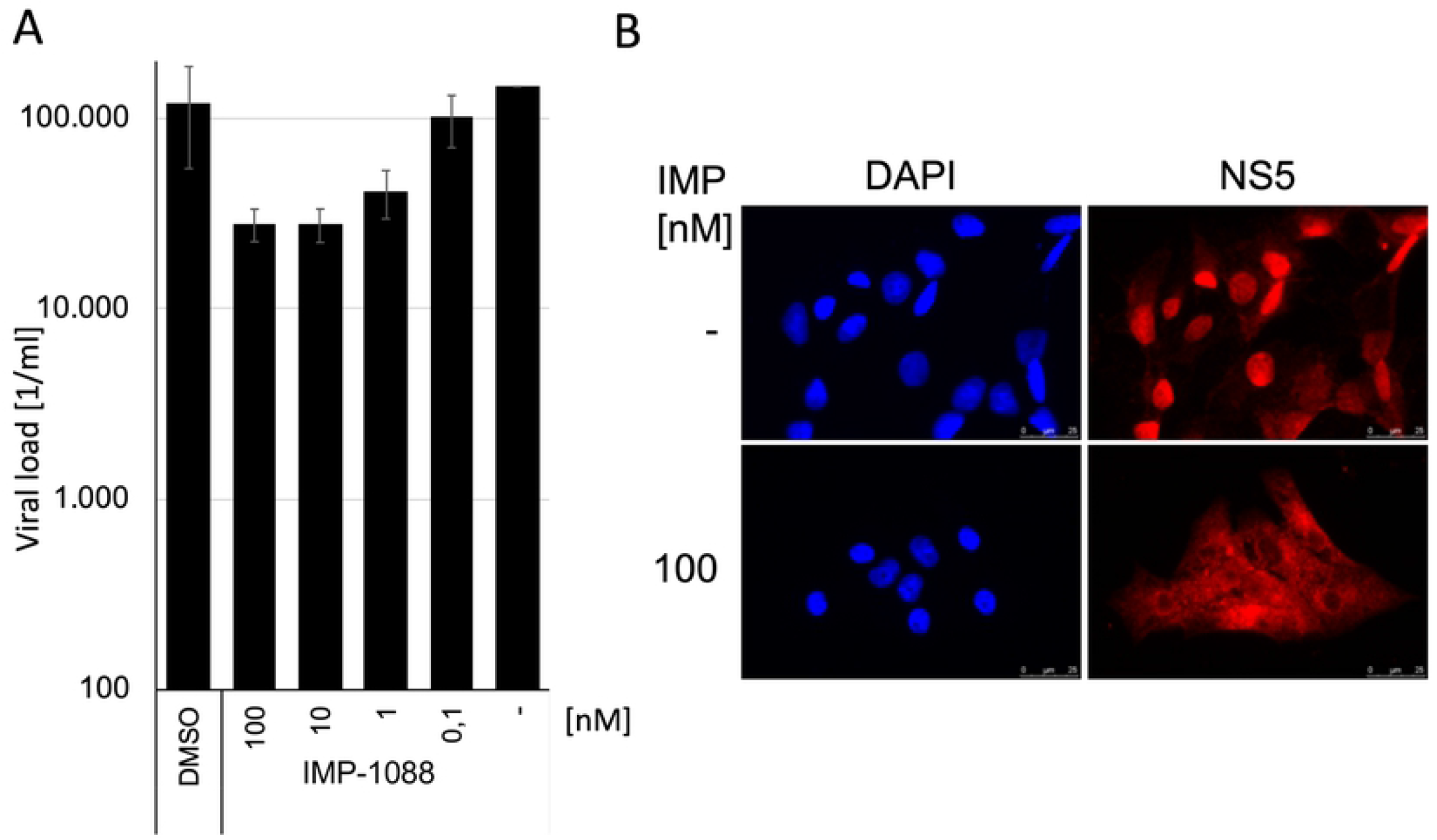
IMP-1088 suppresses Dengue virus replication and influences the subcellular localisation of NS5. **(A)** Vero cells were incubated with IMP-1088 and infected with the Dengue virus in triplicates. Bars indicate the mean of three independent infections. The error bars represent the standard deviation. **(B)** The subcellular localisation of Dengue NS5 was determined by immunofluorescence with a Dengue virus NS5 specific antibody.

Previously it was shown that the subcellular localisation of the DENV2-NS5 protein is controlled by a short stretch of amino acids at the C-terminus and that NS5 shuttles between the cytoplasm and the nucleus. This prompted us to analyse whether the IMP-1088 would influence the balance of the subcellular localisation of DENV2 NS5. Vero cells were seeded on coverslips and infected with DENV2. Two days after infection, cells were fixed and permeabilised. Immunofluorescence with anti-NS5-antibodies confirmed the nuclear localisation of DENV-2 NS5 (Figure 4). However, treatment with IMP-1088 resulted in a redistribution of the NS5 protein to the cytoplasm (Figure 4), indicating that a membrane-binding is a prerequisite of nuclear localisation.

### Zika, Rubella, Chikungunya, and Enterovirus 71 viruses are insensitive to IMP-1088

Since Polioviruses, DENV, and YF viruses responded to IMP-1088, we analysed whether IMP-1088 is active against other tropical viruses such as Zika and Chikungunya virus. Comparing the Zika genome sequence to the YF NS5 sequences revealed that the aminoterminal glycin residue at the NS5 cleavage site is missing in the Zika sequences. This prevents NS5 myristoylation. Furthermore, the processing of all other cleavage sites did not result in the generation of predictable myristoylation sites. Thus, the Zika virus represents a control for unspecific IMP-1088 side effects. PBMCs were treated with IMP-1088 and infected with the Zika virus. The cell culture supernatants were collected two days after infection, and RTqPCR determined viral genome amounts. IMP-1088 did not influence Zika virus replication, indicating that the inhibition of the myristoylation of viral proteins and not cellular factors are the target of IMP-1088 (Figure 5). The experiments were reproduced in Vero cells with similar results (data not shown).

**Figure 5.**
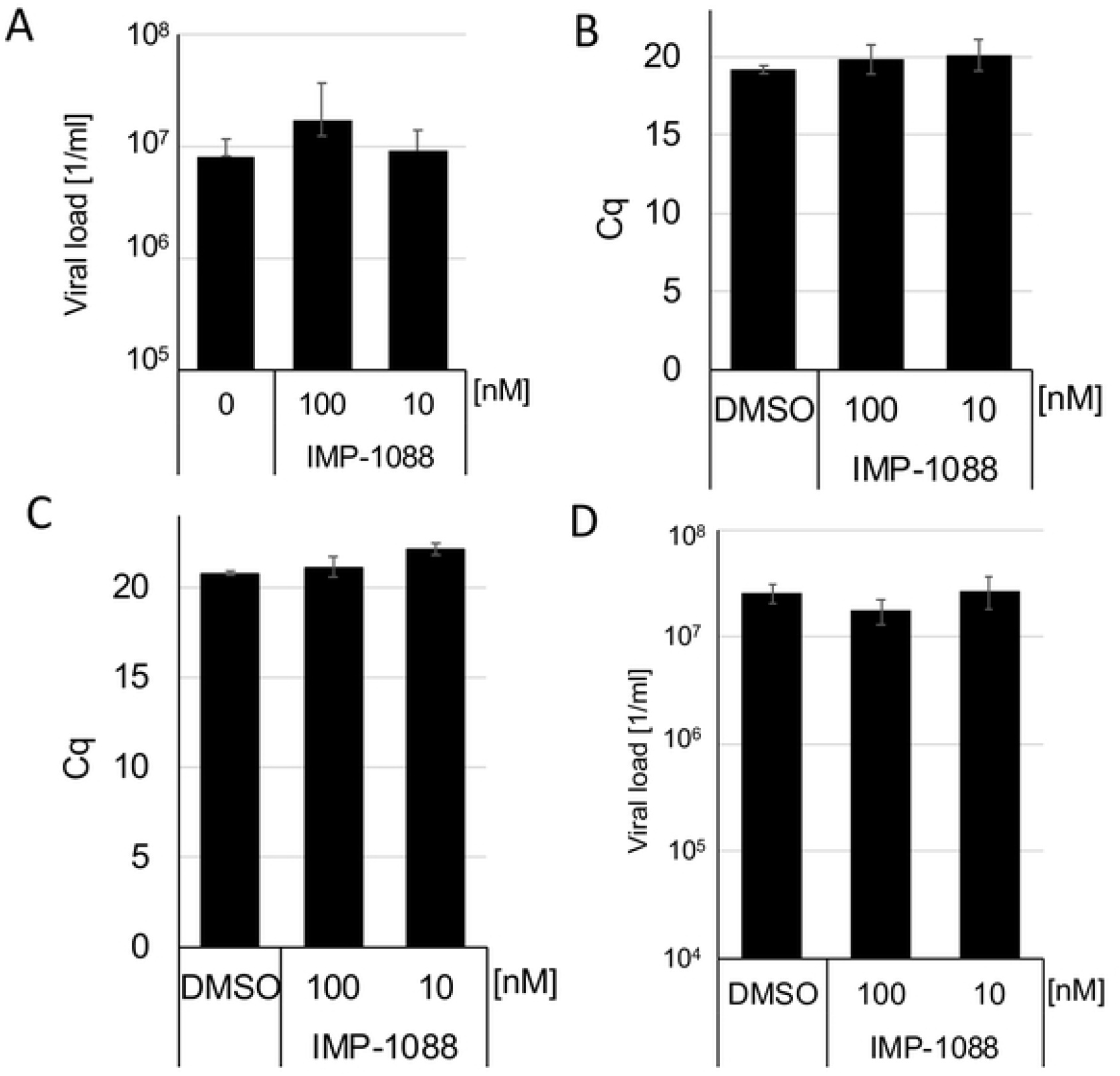
Zika, Chikungunya, Rubella, and Enterovirus 71 are insensitive to IMP-1088. **(A)** PBMC were treated with IMP-1088 and infected with the Zika virus. Genome copy numbers were determined by RTqPCR two days after infection. Bars indicated the standard mean of three independent infections. **(B)** Cell culture cells were incubated with IMP-1088 and infected with either Chikungunya **(B)**, Rubella **(C)**, or Enterovirus 71 **(D)**. Viral RNAs were isolated from the supernatants two days after infection, and viral genomes were quantified by RTqPCR.

To further investigate whether IMP-1088 inhibits positive-stranded RNA viruses, which do not encode predicted myristoylation sequences by inactivating cellular pathways, we sought to analyse Chikungunya, Rubella, and Enterovirus 71 replication.

Bioinformatic analyses of the Chikungunya sequence revealed no predictable myristoylation site even after protein maturation. Nevertheless, BHK-21 cells were incubated with IMP-1088 and infected with the Chikungunya virus. Cellular supernatants were collected two days after infection, and RTqPCR quantified viral RNAs. Although Chikungunya is similar to the other plus-stranded RNA viruses, its replication was not significantly influenced by IMP-1088 (Figure 5), which underlines the specificity of the results obtained with HIV, DENV and the YF. Furthermore, we analysed the potential effects of IMP-1088 on Rubella virus, another plus-stranded RNA virus. Vero cells were incubated with IMP-1088 and subsequently infected with the Rubella virus. Cell culture supernatants were harvested two days post-infection, and viral RNAs were isolated using the MagNa Pure 24 system. The genome amounts were quantified with RTqPCR. In contrast to other plus-stranded RNA viruses, IMP-1088 did not influence the replication of the Rubella virus showing the specificity of the described effects again.

Next, we analysed the influence of IMP-1088 on Enterovirus 71 replication. RD cells were treated with IMP-1088 and infected with patient-derived Enterovirus 71. Cell culture supernatants were harvested two days after infection, viral RNAs were isolated, and viral loads are determined by RTqPCR. Again, IMP-1088 did not influence the replication of Enterovirus 71.

The results with Zika, Chikungunya, Rubella and Enterovirus 71 indicate that the suppression of the myristoylation of cellular factors by IMP-1088, at the concentrations used, is not sufficient to inhibit viral replication. Furthermore, treatment of different cell lines and PBMCs did not inhibit viral replication of viruses without myristoylation signals.

### IMP-1088 inhibits DNA viruses such as Vaccinia virus, HHV-8 and mCMV

It has been reported that E7R, G9R, A16L, A47L, L1R proteins of the Vaccinia virus are myristoylated (11-14). This prompted us to analyse whether IMP-1088 would inhibit Vaccinia virus replication. Vero cells were incubated with IMP-1088 and subsequently infected with the *gfp*-encoding Vaccinia virus. *Gfp*-expressing cells were counted two days after infection (Figure 6A). IMP-1088 at concentrations of 10nm and 100nM suppressed *gfp* expression approximately 0.5 and 1.5 orders of magnitude, respectively. These results indicate that myristoylation of Vaccinia proteins is essential for viral gene expression in the infected cell. Since L1R and A16L are part of the fusion complex, we decided to analyse the effect of IMP-1088 on viral infectivity. Thus, we collected the supernatant of the treated and infected Vero cells to determine the influence of IMP-1088 on the viral infectivity. Again, *gfp*-expressing Vero cells were counted, and the viral infectivity was calculated (Figure 6A). However, the viral infectivity was decreased, similar to the infection in the previous experiment. This indicates that myristoylation of L1R and A16L might not be required for infection in cell culture or that the MNT1/2 activity was still sufficient to modify both viral proteins. Nevertheless, the inhibition of the Vaccinia virus adds to the broad range of inhibition of IMP-1088.

**Figure 6.**
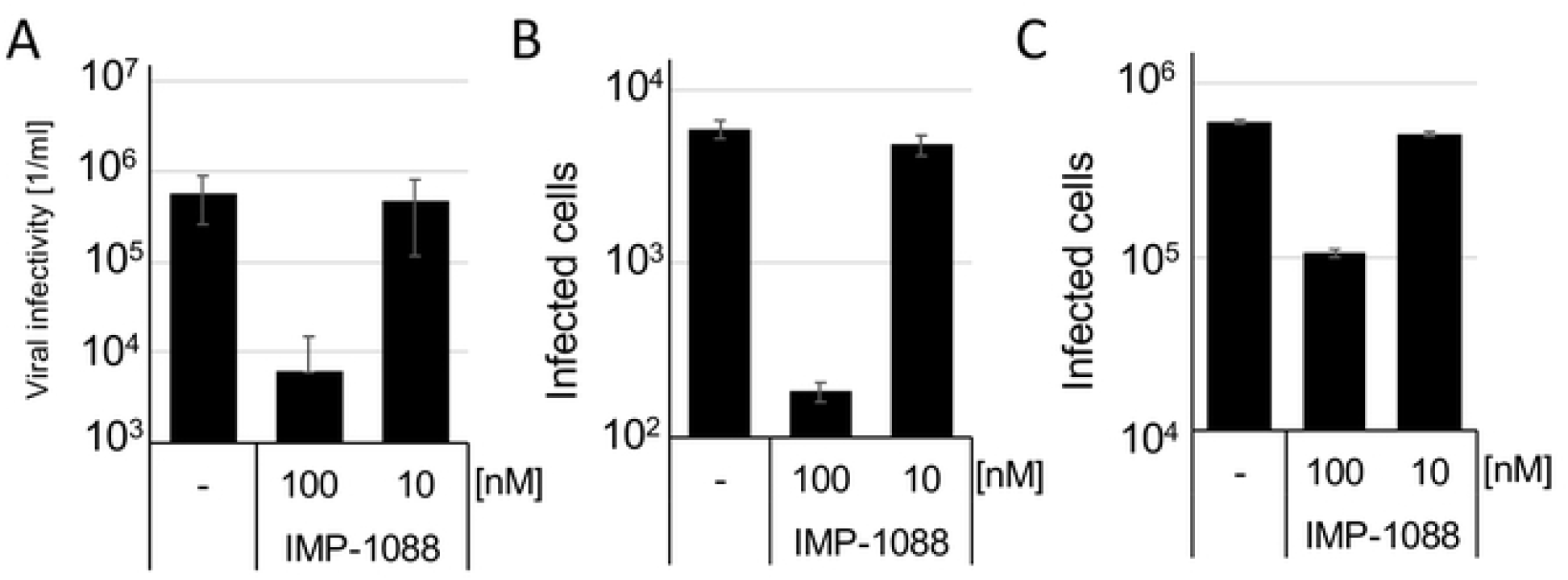
IMP-1088 inhibits Vaccinia virus and Herpesvirus replication. **(A)** Vero cells were treated with IMP-1088 and infected with the *gfp*-encoding Vaccinia virus. GFP expressing cells were counted after one day (left panel). The influence of IMP-1088 on viral infectivity was analysed by infecting cells with the viral supernatants (right panel). **(B)** NIH-3T3 cells were infected with a DsRed encoding murine CMV. Infected cells were counted two days after infection. **(C)** BHK-21 cells were treated with IMP-1088 and subsequently infected with g*fp*-encoding HHV-8. Infected cells were counted two days post-infection.

Analyses of the published Herpesvirus sequences revealed several potential myristoylation proteins (15). Besides, it has been shown that HSV1 represses the myristoylation of cellular proteins. Therefore, the effects of IMP-1088 on Betaherpesviruses (murine CMV) and Gammaherpesviruses (HHV-8) were analysed.

To further investigate whether Herpesviruses require myristoyltransferase activity, NIH-3T3 cells were incubated with IMP-1088 and subsequently infected with murine CMV (mCMV). The mCMV virus used expresses DsRed under the control of a late promoter. The entry in the *late*-phase indicated by DsRed expression was suppressed by one order of magnitude (Figure 6B). These results provide evidence that both the *early*-gene expression and the entry into the *late*-phase of mCMV need the myristoyltransferase activity.

Finally, we determined whether HHV-8 is susceptible to IMP-1088. BHK-21 cells were incubated with IMP-1088 and subsequently infected with *gfp*-encoding HHV-8. IMP-1088 at concentrations of 100nM decrease *gfp*-expressing cells about half an order of magnitude after 48h, while virus spread after 72h was suppressed by one log-step (Figure 6C). This indicates that HHV8 is inhibited by IMP-1088, which reduces influences, especially the virus-spread in cell culture.

In summary, we provide evidence that Vaccinia virus, mCMV and HHV-8 need cellular myristoyltransferase for optimal replication in cell culture. However, the effect of IMP-1088 is less pronounced compared to the plus-stranded RNA viruses.

In general, usually, antiviral therapies are virus-specific and target viral enzymes. This sets viruses on an evolutionary pressure and leads to the selection of viral escape mutants. The resistances associated mutants have been extensively studied for HIV-1, where in some cases, more than 20% of the PR encoding amino acid residues were exchanged during antiviral therapy. Targeting of the cellular myristoyltransferase 1 and 2 would block in case of HIV two essential steps, Gag assembly at the plasma membrane and Nef function. It is unlikely how the virus could adapt since the functions of two critical protein are affected. In this regard, IMP-1088 is different from previously used antivirals which inhibited single viral proteins. Furthermore, the inclusion of IMP-1088 in combination therapies would provide an additional treatment option for the patient all therapeutic options exhausted and for HHV-8 seropositive HIV-1 infected patients.

The Flaviviral NS5 protein is highly conserved and multifunction. Here, we presented evidence that YF NS5 is myristoylated and that myristoylation is required for viral infectivity of Dengue and Yellow fever viruses. This is supported by the IMP-1088 resistance of Zika viruses, which lack any potential myristoylation site in its sequence. This result indicates that the inhibition of myristoylation of viral DENV and YF NS5 proteins is responsible for the antiviral effect of IMP-1088. Similar results were obtained in previous experiments with myristoyl transferase 1-knockdown cells showing that NMT activity is required for DENV replication (16). The negative impact on other RNA viruses provides evidence that the observed effect is virus-specific, not related to the inhibition of general cellular factors.

Flaviviruses replicate in the cytoplasm. Thus, it is likely that viral proteins are localised there. However, previous reports and we described localisation of the YF NS5 protein in both the nucleus and the cytoplasm (17). The inhibition of the myristoylation with IMP-1088 resulted in almost exclusive nuclear localisation, indicating that the cytoplasmic YF NS5 might interact with cellular membranes. In contrast, the NS5 protein of the DENV Serotype 2 encodes functional nucleolar localisation signals and is almost exclusively localised in the nucleus (18, 19). Treating cells with IMP-1088 led to the nuclear exclusion of DENV-2 NS5 protein. The nucleolar localisation contributes to the dysregulation of the host-translation and splicing, which enhances virus replication, and importin inhibitors led to reduced virus replication (20, 21). This could explain the observed reduction of DENV-2 titers in IMP-1088 treated cells.

In summary, inhibition of the cellular myristoyltransferase can specifically and efficiently suppress the replication of RNA and DNA viruses.

## Methods

### Viruses, viral proteins and chemicals

Yellow fever vaccine strain: STAMARIL 17D-204; Poliovirus Serotype 3 vaccine strain; Dengue virus type 2 strain (22), Chikungunya strain Würzburg (23), Zika virus plasmids pH/PF/2013 and pMR766/2013 were kindly provided by R. Bartenschlager (Heidelberg). The Yellow fever NS5 protein was codon-optimised using the algorithms provided by Invitrogen (Thermo Fisher, USA), and a FLAG-tag encoding sequence was added to the carboxyl-terminus. The synthetic DNA was cloned into the pcDNA3.4 vector. Enterovirus-71 (RD cells) and Rubella virus (Vero cells) were patient-derived isolates obtained from the diagnostics of the department of the virology Würzburg and were used with permission. HIV-1 experiments were performed with the pNL4-3-derived virus (24). All infections were performed in at least triplicate assays and repeated three times. IMP-1088 was purchased from Cayman Chemical, USA.

### Cellular proliferation assays

The proliferation of cells with and without IMP-1088 was determined by direct automatic cell counting. Cells were seeded on optical plates (CellCarrier-96, PerkinElmer) and counted before the experiments. Then IMP-1088 was added in decreasing concentrations, and the cells were incubated for three days. The cell numbers per well were determined using the PerkinElmer Ensight reader. Only compound concentrations that did not reduce the cell number per well significantly were used for antiviral assays.

### HIV-1 infectivity assay

Viral infectivity was determined using TZM-bl indicator cells as described before (25). In brief: PBMCs were infected with pNL4-3 derived HIV-1 viruses. Supernatants of the infected cells were centrifuged at 2000rpm for 5min to remove all detached cells and titrated on indicator TZM-bl cells. TZM-bl cells are CD4+, CCR5+, and CXCR4+ HeLa cells, which encode an integrated HIV-LTR controlling the expression of a beta-Galactosidase. After two days, the cells were fixed with methanol/acetone (1:1), extensively washed with PBS and ß-Galactosidase activity was visualised with X-Gal substrate. The blue-stained infected cells were counted, and viral titers were calculated.

### Viral RNA isolation and RTqPCR

Viral RNAs were isolated with the viral NA isolation kit (Roche, Germany) or with the MagNa Pure 24 NA isolation device according to the manufacturer’s instructions (Roche). Genomes quantification of YF was done as described before (10). All other viruses were quantified with LightMix Assays (Roche). The RNA Master Probe kit was used for amplification as described by the manufacturer (Roche). PCR-Setup was performed with the Liquid Handling Station (BRAND, Germany) in triplicate assays. All PCRs were performed using either the LightCycler96 or 480II (Roche). Quantifications were performed with the respective cycler software. EC50 values were calculated using GraphPad Prism.

### Immunofluorescence and Transfection

Vero cells were seeded on coverslips, incubated with IMP-1088 and infected with DENV2 or YF. Cells were fixed with 5% PFA and permeabilised with 0.25% Triton X in PBS. Unspecific binding of the antibodies was blocked with 3% bovine serum albumin. Viral NS5 proteins were visualised with an NS5 specific antiserum (Biozol, Germany) and a Cy3 coupled donkey anti-rabbit IgG antibody (Dianova, Germany). HepG2 cells were transfected with the pcDNA3.4-synNS5-FLAG. The localisation of the codon-optimised NS5 protein was detected with a rabbit-anti-FLAG antibody (Sigma, Germany).

## Acknowledgement

We would like to thank the viral diagnostics of the department of Virology Würzburg for patient-derived virus isolates and BRAND for providing the LHS.

